# Frequency- and area-specific entrainment of intrinsic cortical oscillations by repetitive transcranial magnetic stimulation

**DOI:** 10.1101/455857

**Authors:** Yuka O. Okazaki, Yumi Nakagawa, Yuji Mizuno, Takashi Hanakawa, Keiichi Kitajo

## Abstract

Neural oscillations are ubiquitous throughout the cortex, but the frequency of oscillations differs from area to area. To elucidate the mechanistic architectures establishing various rhythmic activities, we tested whether spontaneous neural oscillations in different cortical modules can be entrained by direct perturbation with distinct frequencies of transcranial magnetic stimulation (TMS). While recording the electroencephalogram (EEG), we applied single-pulse TMS (sTMS) and repetitive TMS (rTMS) at 5, 11, and 23 Hz to motor or visual cortex. To assess entrainment, defined as phase locking of intrinsic oscillations to periodic external force , we examined local and global modulation of the phase-locking factor (PLF). sTMS triggered transient phase locking in a wide frequency band with distinct PLF peaks at 21 Hz in the motor cortex and 8 Hz in the visual cortex. With TMS pulse trains of 11 Hz over visual cortex and 23 Hz over motor cortex, phase locking was progressively enhanced at the stimulation frequency and lasted for a few cycles after the stimulation terminated. Moreover, such local entrainment propagated to other cortical regions, suggesting that rTMS entrained intrinsic neural oscillations locally and globally via network nodes. Because the entrainment was frequency-specific for each target site, these frequencies may correspond to the natural frequency of each cortical module and of the global networks. rTMS enables direct manipulation of the brain and is thus useful for investigating the causal roles of synchronous neural oscillations and synchrony in brain functions, and for the treatment of clinical symptoms associated with impaired oscillations and synchrony.

**Significance Statement:** We provide the first evidence for area- and frequency-specific entrainment by frequency-tuned repetitive transcranial magnetic stimulation (rTMS), and the propagation of this entrainment to other areas. Our results indicate that rTMS at the natural frequency of each cortical system is particularly effective for entraining oscillatory phase. Moreover, local entrainment led to global entrainment in functionally coupled areas. The ability to control brain rhythms in the intact human brain is highly beneficial for studying the causal roles of rhythmic activity in brain function. Moreover, this modulatory technique has the potential to treat patients with impaired rhythmic networks in disorders such as schizophrenia and stroke.

## 1. Introduction

Accumulating evidence suggests that neural oscillations play a functional role in the brain. Although neural oscillations are ubiquitous in the brain, oscillation frequency varies across distinct brain regions and networks. For example, oscillations from 8 to 13 Hz are typically observed in occipital visual areas (alpha rhythm) and over parietal sensorimotor areas (mu rhythm). In the sensorimotor areas, oscillations at higher frequencies (13 to 30 Hz; beta rhythm) are also observed, but the mu and beta rhythms have different sources, originating from the somatosensory cortex and the motor cortex, respectively (Hari and Salmelin, 1997; Okazaki et al., 2014). From a global network perspective, frequency-specific, large-scale phase synchronization of oscillations is important in linking task-relevant brain regions associated with information processing (e.g., face perception, Rodriguez et al. (1999); selective attention, Osipova et al. (2008); working memory, Kawasaki et al. (2010)). Thus area-specific oscillations with distinct frequencies presumably help establish functional networks.

Frequency-specific spatial structures established by phase synchronization, such as the alpha- and beta-band synchronization in the visual and sensorimotor networks, also exist in the resting state (Hillebrand et al., 2012; Hipp et al., 2012) and seem to be spatially consistent with the task-driven networks, as demonstrated by fMRI studies (Ergenoglu et al., 2004; Kawasaki et al., 2014a). Moreover, research in animals has shown that sensory-evoked population firing patterns are geometrically confined to subregions of the neuronal state space delineated by spontaneous activity (Luczak et al., 2009). These studies suggest that the spatiotemporal patterns of spontaneous neural activity reflect the full repertoire of neural dynamics, and constrain the task-related neural dynamics. However, few mechanistic details are known about the various sets of rhythmic activity in different cortical modules.

Noninvasive brain-stimulation techniques such as transcranial magnetic and electrical stimulation (TMS and tES) have emerged as promising manipulative tools, enabling direct perturbation of local brain modules. TMS, in particular, can be used to target spatially confined regions involved in generating oscillations. Moreover, single-pulse TMS can directly interfere with the phase dynamics of oscillations, inducing phase resetting of intrinsic oscillations in visual areas (Kawasaki et al., 2014b). Single-pulse TMS over the motor (Paus et al., 2001) and visual cortices (Herring et al., 2015) can also trigger intrinsic cortical oscillations in the beta- and alpha-bands, respectively. It is noteworthy that when single-pulse TMS was applied to different thalamocortical regions that constitute specific networks, oscillations induced in occipital, parietal, and frontal cortices fell into distinct frequency bands (Rosanova et al., 2009). This indicates that each thalamocortical module has its own characteristic intrinsic frequency, or “natural frequency”. Given the results of single pulse-induced brain oscillations, entrainment of oscillations can be predicted when further TMS pulses are applied in phase with the induced oscillations. Consequently, the amplitude of oscillations gradually increases as more and more intrinsic neural oscillators are entrained to the repetitive TMS pulses (Kayser and Tenke, 2006; Thut et al., 2011a). In line with this hypothesis, Thut et al. (2011b) demonstrated local enhancement of alpha oscillations by applying alpha-frequency trains of TMS pulses to the parietal cortex.

However, to our knowledge, no previous study has applied rhythmic stimulation at multiple frequencies to distinct cortical regions; thus it remains unclear whether periodic stimulation results in entrainment with different frequency characteristics for each local region. Moreover, it is not clear how locally entrained oscillations impact oscillations in other cortical regions. Given that local brain modules that are targeted by periodic stimulation are globally coupled to oscillatory modules in other brain regions, we predict that the entrained oscillations will propagate to other connected areas with specific frequency characteristics. To address these questions, we measured phase dynamics with electroencephalography (EEG) while applying rTMS to either the motor or visual cortex at theta- (5 Hz), alpha- (11 Hz), or beta- (23 Hz) band frequencies.

## 2. Materials and Methods

### 2.1 Participants

Fourteen healthy right-handed participants (two female and twelve male, aged 30.8 ± 5.5 years, mean ± s.d.) gave informed, written consent to participate in the study. This study was approved by the RIKEN ethical committee.

### 2.2 TMS–EEG experiments

#### TMS

TMS was applied in a biphasic pulse configuration by a Magstim Rapid unit with a figure-of-eight coil (Double 70 mm Alpha coil; Magstim, UK). The stimulation site was over either visual or motor regions, or a sham control location (see Fig. 1A). For sham stimulation, the coil was positioned 10 cm above the vertex with the coil handle directed posterior. Thus, sham stimulation produced the TMS “click” sound but without cortical stimulation; in all conditions, the click sound was attenuated by earplugs. For stimulation of visual regions, the coil was located centrally between the Oz and O2 electrodes with the coil handle pointing rightward and the coil surface parallel to the surface of the scalp. For stimulation of motor regions, the coil position was determined individually at the “hotspot” that activated the right first dorsal interosseous (FDI) (approximately at the C3 electrode with the handle perpendicular to the central sulcus). A stimulation intensity of 90% of the FDI active motor threshold was used in all conditions.

**Figure 1.**
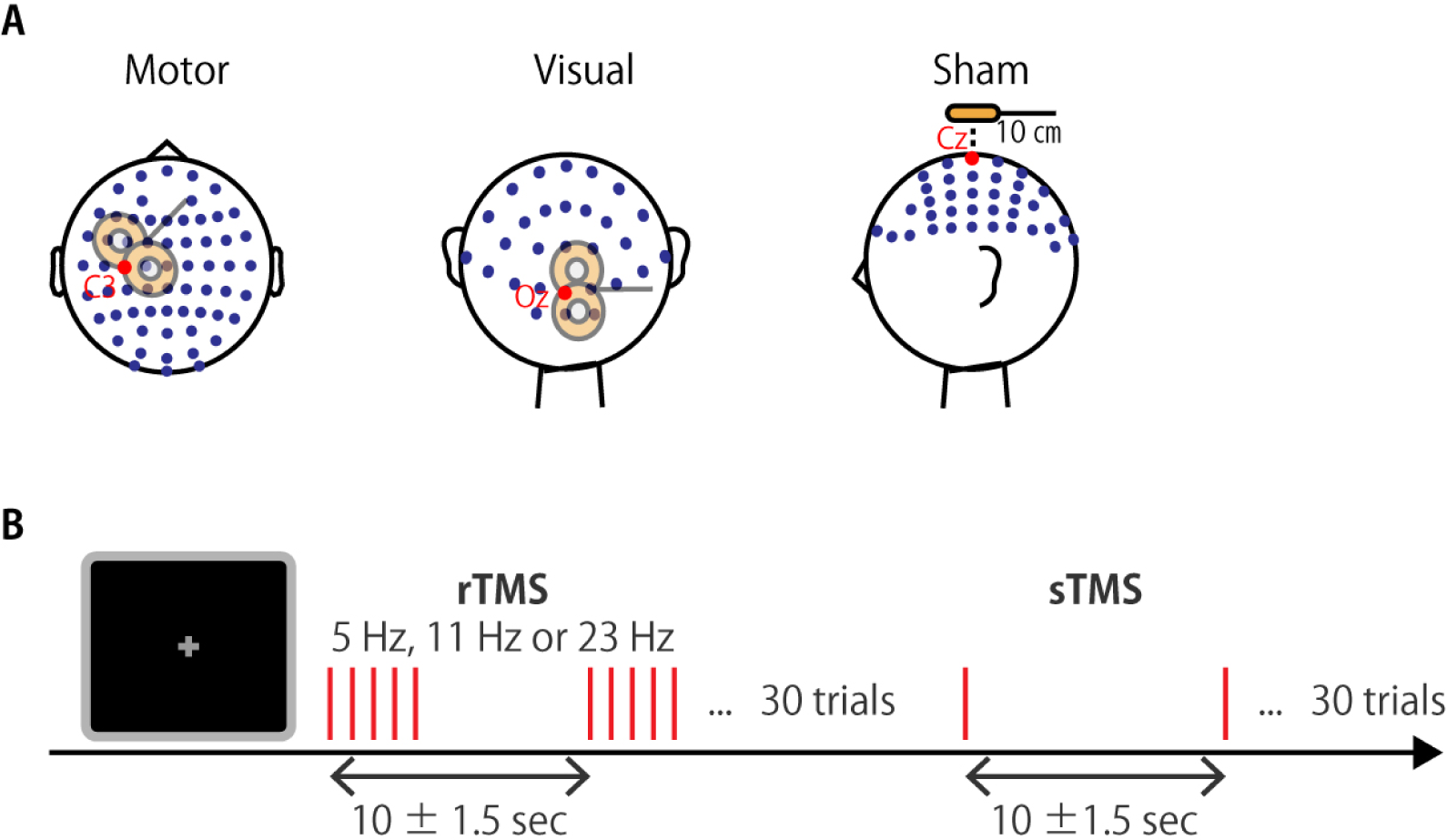
Experimental paradigm. (A) Schematic illustration of the stimulation sites. (B) The TMS conditions were a single pulse (sTMS), and five pulses at 5 Hz, 11 Hz, and 23 Hz (rTMS). Conditions were presented in blocks with 30 repetitions. Participants had to fixate a central cross during trials.

##### EEG recordings

During stimulation, EEG (left earlobe reference; ground AFZ) was continuously recorded at a 5 kHz sampling rate (filtering: DC to 1000 Hz) from 63 scalp sites via Ag/AgCl TMS-compatible electrodes mounted on 10/10 system EasyCap (EASYCAP GmbH, Germany). Horizontal and vertical electrooculography (EOG, ground electrode was placed on the left mastoid) were similarly continuously recorded at a 5 kHz sampling rate. Electrode impedance was maintained below 10 kΩ. The EEG and EOG signals were amplified by a Brain Amp MR+ system (Brain Products GmbH, Germany).

##### Procedure

Three TMS–EEG sessions with different stimulation sites (visual, motor, and sham) were conducted in a random order. Each session consisted of four blocks with different pulse trains in each block: a single pulse (sTMS), and 5-pulse trains at 5 Hz (θ-rTMS), 11 Hz (α-rTMS), and 23 Hz (β-rTMS). Each block contained 30 trains with an inter-train interval of 10 ± 1.5 s. Participants were fixated on a centrally presented gray cross on a black background during each block (Fig. 1B). Stimulus delivery was controlled by MATLAB (Mathworks, USA) with the Psychtoolbox-3 extension (Brainard, 1997; Pelli, 1997; Kleiner, 2007). The TMS-EEG data were also analyzed in our study for a different purpose, namely to probe phase-amplitude coupling (Glim, 2017).

#### 2.3 Preprocessing

EEG data were analyzed with custom MATLAB (Mathworks, Natick, MA, USA) scripts developed in-house, with FieldTrip (Oostenveld et al., 2011) and EEGLAB (Delorme and Makeig, 2004), open source MATLAB toolboxes for the analysis of neurophysiological data. First, EEG data were segmented from 2 s before the first pulse to 3 s after the last pulse. Then, the segmented epochs were re-referenced offline to the average of the left and right earlobe electrode signals.

TMS artifacts and noisy epochs were rejected by performing the following steps according to the study by Herring et al. (2015) (see also http://www.fieldtriptoolbox.org/tutorial/tms-eeg for the detailed procedure). First, we linearly interpolated the 40 samples (8 ms) after TMS onset, which is the period that usually shows excessive TMS artifacts. In the case of artifacts occurring after 8 ms, we used interpolation of 60 samples (12 ms). Second, we attenuated the exponentially decaying TMS artifacts using independent component analysis (ICA) (Korhonen et al., 2011). Independent components (ICs) with extremely large amplitudes, i.e., those with maximum z-score values greater than 1.65 between 0 and 100 ms, were removed. Third, we discarded epochs within 1 s pre- or post-stimulation in which the EEG value exceeded 200 μV. Then, we applied current source density (CSD) transformation to the EEG voltage map using the spherical-spline surface Laplace algorithm (Perrin et al., 1989; Kayser and Tenke, 2006) to attenuate the effects of volume conduction using CSD Toolbox (http://psychophysiology.cpmc.columbia.edu/Software/CSDtoolbox). Finally, the data were downsampled to 1000 Hz.

#### 2.4 Time–frequency analysis

We obtained time–frequency representations (TFRs) of the instantaneous amplitude and phase using a wavelet transform. Convolution of the EEG signal *x(t)* with a Morlet wavelet 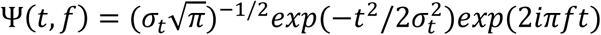 gives a complex matrix *W(t, f)*, of which the square norm corresponds to the amplitude and the arctangent corresponds to the phase of the signal *x(t)* at a center frequency *f* and time *t*, with standard deviations *σ_f_ = 4f/m* and *σ_t_ = m/2πf* (Tallon-Baudry et al., 1996; Lachaux et al., 2000). The constant *m*, which defines the time and frequency resolution of the trade-off to control for bias due to the different number of trials that survived between conditions (Fisher, 1993). We also confirmed that ZPLF remains flat with increasing number of relationship, was set to 3.

Entrainment is defined as the phase locking of ongoing neuronal oscillations to a periodic external force. Thus, it is expected that phase relations over trials become consistent through phase locking to frequency-tuned rhythmic stimulation. To assess this, we computed the phase-locking factor (PLF) (Tallon-Baudry et al., 1996):

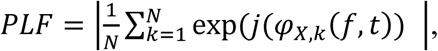

where φ_x_ is the instantaneous phase of frequency *f* at electrode channel *X* and *N* denotes the number of trials. In general, the PLF of a small number of trials was higher than that of a high number of trials. Supplemental Figure S1 shows PLF as a function of the number of randomly chosen trials from a pool of trials under all stimulated conditions from a single subject. Because the bias caused by a small number of trials was no longer observed at about 10 trials (see Supplemental Information: Fig. S1A), the number of trials in our experiments (mean±sd, 24.8±2.6 trials) was above the minimal number of trials required to remove this bias. Moreover, the PLF for each participant was Rayleigh Z-transformed

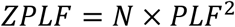

to control for bias due to the different number of trials that survived between conditions (Fisher, 1993). We also confirmed that ZPLF remains flat with increasing number of trials (see Supplemental Information; Fig. S1B).

#### 2.5 Statistical analyses

Significant differences in ZPLF between TMS (sTMS and rTMS) and sham TMS were determined by cluster-based permutation tests (Maris and Oostenveld, 2007), which resolve the problem of multiple comparisons over channels (63 channels), frequencies (3–45 Hz), and time points (−0.5–1.5 s). First, we compared every point (i.e., channel, frequency, time) in the ZPLF matrices for the TMS and sham-TMS conditions using a two-tailed paired *t*-test with a threshold uncorrected *p*-value < 0.05 to locate contiguous negative and positive clusters in the matrices. Cluster size was assessed as the sum of the *t*-value within the cluster. Second, to generate a null distribution for the cluster-level test statistics, we identified the largest cluster from matrices in which two condition labels were randomly permuted within participants and iterated 500 times. We defined the 97.5th percentile of the null distribution as the level for statistical significance and used this to identify significant clusters in the observed data.

To assess whether the modulation of ZPLF is more globally distributed at frequencies matching the stimulation frequency than at other frequencies, we first identified channels with significant ZPLF using the cluster-based permutation test described above. We then counted the number of significant channels at each frequency and time. Finally, we compared the number of significant channels at the stimulation frequency and at other frequencies using the binomial test. Statistical significance was set at *p* < 0.05.

## 3. Results

To demonstrate the effects of sTMS and rTMS at theta-, alpha-, and beta-band stimulation frequencies on the phase dynamics of oscil latory neural activity, we examined phase consistency in stimulation trials compared with sham trials, where the phase relationship should be close to random.

### 3.1 Entrainment of intrinsic local oscillations by periodic stimulation

To assess the entrainment of neural oscillations by TMS, we first examined the ZPLF as time–frequency representations (TFR) for each stimulated area (Fig. 2). The cluster-based permutation test revealed significant increases in phase locking by sTMS and rTMS (*θ-*rTMS, *α*-rstimulation-frequency cycles). One-way TMS, *β*-rTMS) compared with the corresponding sham condition (*p* < 0.05, cluster-based permutation test). For stimulation over motor areas, sTMS induced a transient increase in the ZPLF at a broad frequency band, peaking at 21 Hz (Fig. 2A). rTMS induced a continuous increase in the ZPLF at the stimulation frequency (horizontal dashed line) during *α*-rTMS (Fig. 2C) and *β*-rTMS (Fig. 2D), but the stimulation-frequency response was negligible during *θ-*rTMS (Fig. 2B). For stimulation over visual areas, the peak response frequency to sTMS was 8 Hz (Fig. 2E). In all rTMS conditions, rTMS resulted in a continuous increase in entrainment around the stimulation frequency (Fig. 2F-H).

**Figure 2.**
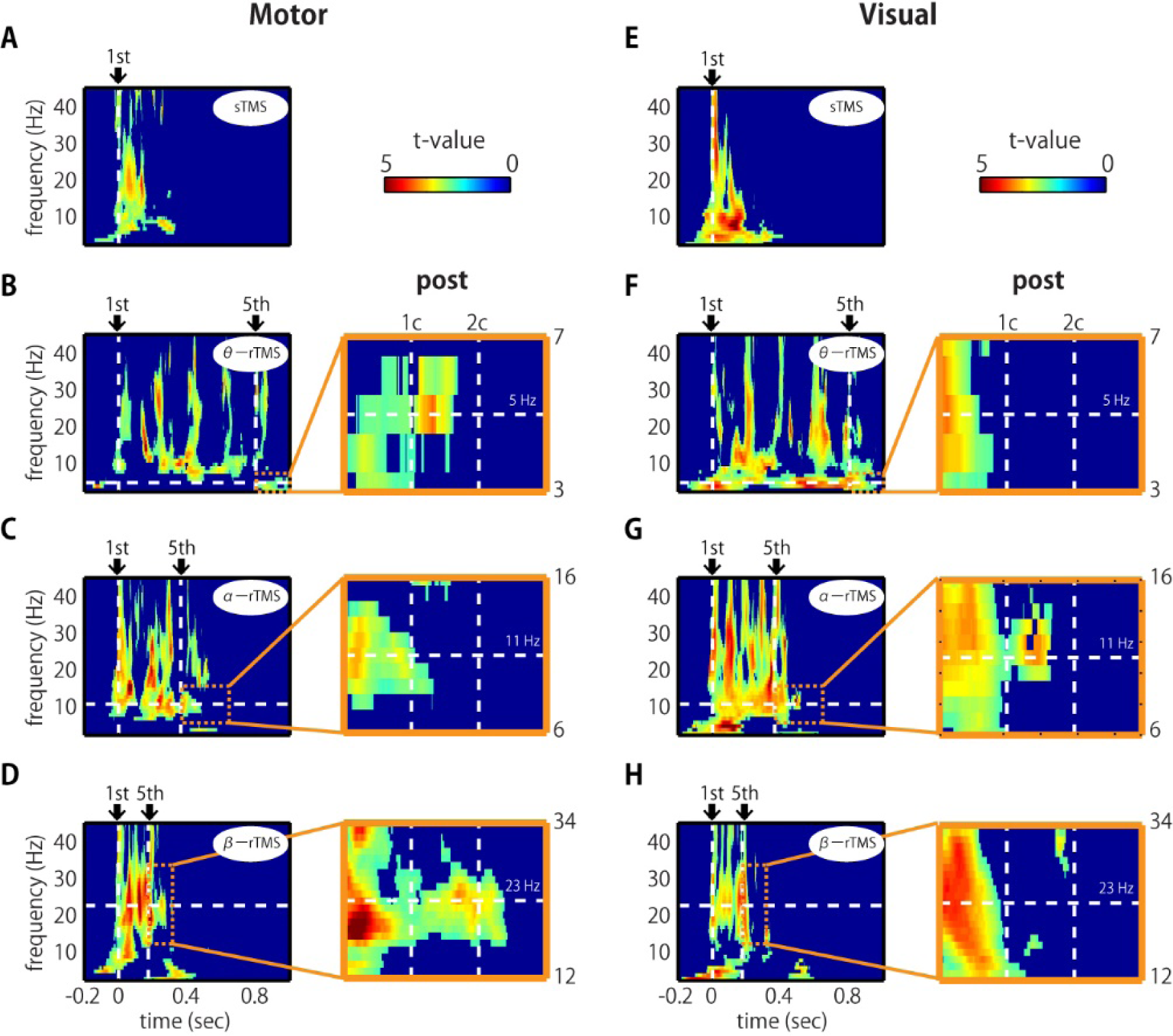
Time–frequency representations of ZPLF *t*-values at the stimulated areas. (A-H) Significantly clustered *t*-values (*p* < 0.05) from the comparisons of ZPLF between motor (C3) and sham stimulation (A-D), and between visual (Oz) and sham stimulation (E-H). The vertical lines in the left panels correspond to first and last stimulation pulses. The horizontal lines indicate the stimulation frequency. Right panels are magnified views of three cycles after the end of the rTMS frequency. Vertical lines indicate post-one cycle (1c) and post-two cycles (2c) for each stimulation frequency.

Next, we asked whether the effects of phase locking across trials last even after the rTMS train has terminated. If intrinsic oscillations are generated by self-sustaining systems without external input (Pikovsky et al., 2003), the phase locking should persist for a short time after the end of the stimulation train (Klimesch et al., 2004). We observed that the increase in ZPLF lasted for more than 2 cycles after the last pulse of β-rTMS to the motor cortex (Fig. 2D, magnified view) and 1.5 cycles after the last pulse of α-rTMS to the visual cortex (Fig. 2G, magnified view). We also note that these sustained frequencies were different from the frequency of oscillations evoked by sTMS. Furthermore, participants whose intrinsic alpha frequency was closer to the stimulation frequency (i.e., 11 Hz) showed a longer entrainment effect (Supp. Fig. 2). These results indicate that rTMS can entrain ongoing neural oscillations in local brain modules.

### 3.2 Gradual modulation of phase-locked frequency over TMS pulses

If entrainment is achieved through a phase alignment of ongoing oscillations to periodic force, successive phase alignment should result in gradual increases in ZPLF (Thut et al., 2011a). As shown in Figure 2, increases in ZPLF occurred not only around the stimulation frequency but also at a broad range of frequencies. Intriguingly, however, the most prominent phase-locked frequency varied over the course of the stimulation train. To measure which frequency showed the most prominent increase at each TMS pulse, we standardized the ZPLF using the mean and SD of the ZPLF over all frequencies (ZPLF_norm_) (Fig. 3; vertical dashed lines indicate the stimulation frequency in each condition). For example, for α-rTMS over the visual cortex, the dominant frequency was 5 Hz at the first pulse (brown line) but 11 Hz for the last three pulses (Fig. 3E). The charts inset in each panel quantify, for each pulse in the train, whether the ZPLF_norm_ at the stimulation frequency was significantly larger than the ZPLFnorm at baseline (the period from −5 to −2 stimulation-frequency cycles). One-way ANOVA showed statistical significance in β-rTMS over the motor cortex (*F*(5, 78) = 3.40, *p* = 0.008) but not in stimulation over the visual cortex (*F*(5, 78) = 1.65, *p* = 0.156). On the other hand, stimulation over the visual cortex showed statistical significance in α-rTMS (*F*(5, 78) = 2.81, *p* = 0.022) and θ-rTMS (*F*(5, 78) = 3.11, *p* = 0.013) . The subsequent Dunnett *post-hoc* tests comparing each pulse with baseline (pre-stimulation interval: −5 - −2 stimulation-frequency cycles) confirmed the gradual increase in phase locking for each pulse train of β-rTMS over the motor cortex (Fig. 3C) and α-rTMS over the visual cortex (Fig. 3E). Although there was a significant difference in θ-rTMS over the visual cortex (Fig. 3D), the phase locking to the stimulation frequency did not increase gradually over the pulse train but was dominant for every TMS pulse in the train.

**Figure 3.**
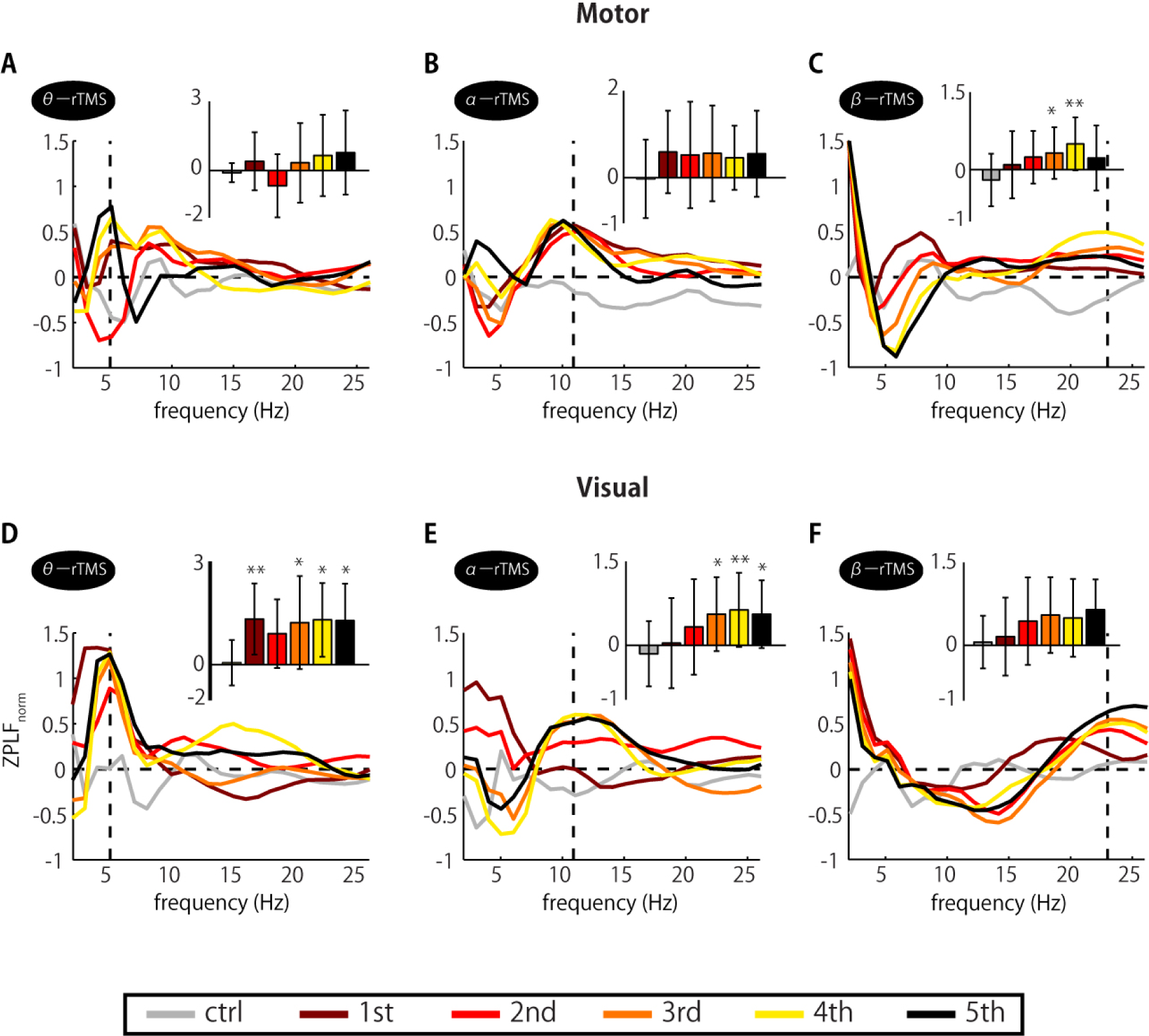
Changes in the dominant frequency of ZPLFnorm (z-transformed ZPLF) during TMS trains. Each colored line corresponds to the response to a particular pulse in the train. Together, they indicate which frequency was the most prominent during the train and how it varied with pulse repetition. Note that each line is the mean value of ±0.5 cycles at each frequency. The charts inset in each panel indicate the ZPLFnorm at the frequency corresponding to the rTMS stimulation frequency, i.e., the values of ZPLFnorm at the vertical dashed line. Significant changes from baseline were assessed by Dunnett’s test (* *p* < 0.05, \*\**p* < 0.01). The baseline was the mean value during the period from −5 to –2 stimulation-frequency cycles. Error bars: STD.

### 3.3 Global propagation of phase entrainment

Next, we addressed the question of whether local phase locking to rTMS propagates globally from the stimulation site. We first examined the spatial extent of significant increases in ZPLF using a cluster-based permutation test on topographical maps of ZPLF. Increases in ZPLF with rTMS to the motor cortex were widely observed in the ipsilateral hemisphere and partially extended to the contralateral hemisphere. For stimulation over visual cortex, rTMS resulted in relatively widespread increases reaching to frontal areas (Fig. 4B). The extensive phase locking during stimulation of the motor and visual cortices appears to be more localized for later pulses within the stimulation train. Moreover, in several areas, phase locking persisted even after the end of the stimulation. In particular, phase locking of frontoparietal areas with θ-rTMS, the left occipitoparietal area with α-rTMS, and the right motor area with β-rTMS were maintained in both visual and motor area TMS-target conditions (see magnified topography at post 1.5 cycles in Fig. 4A and B).

**Figure 4.**
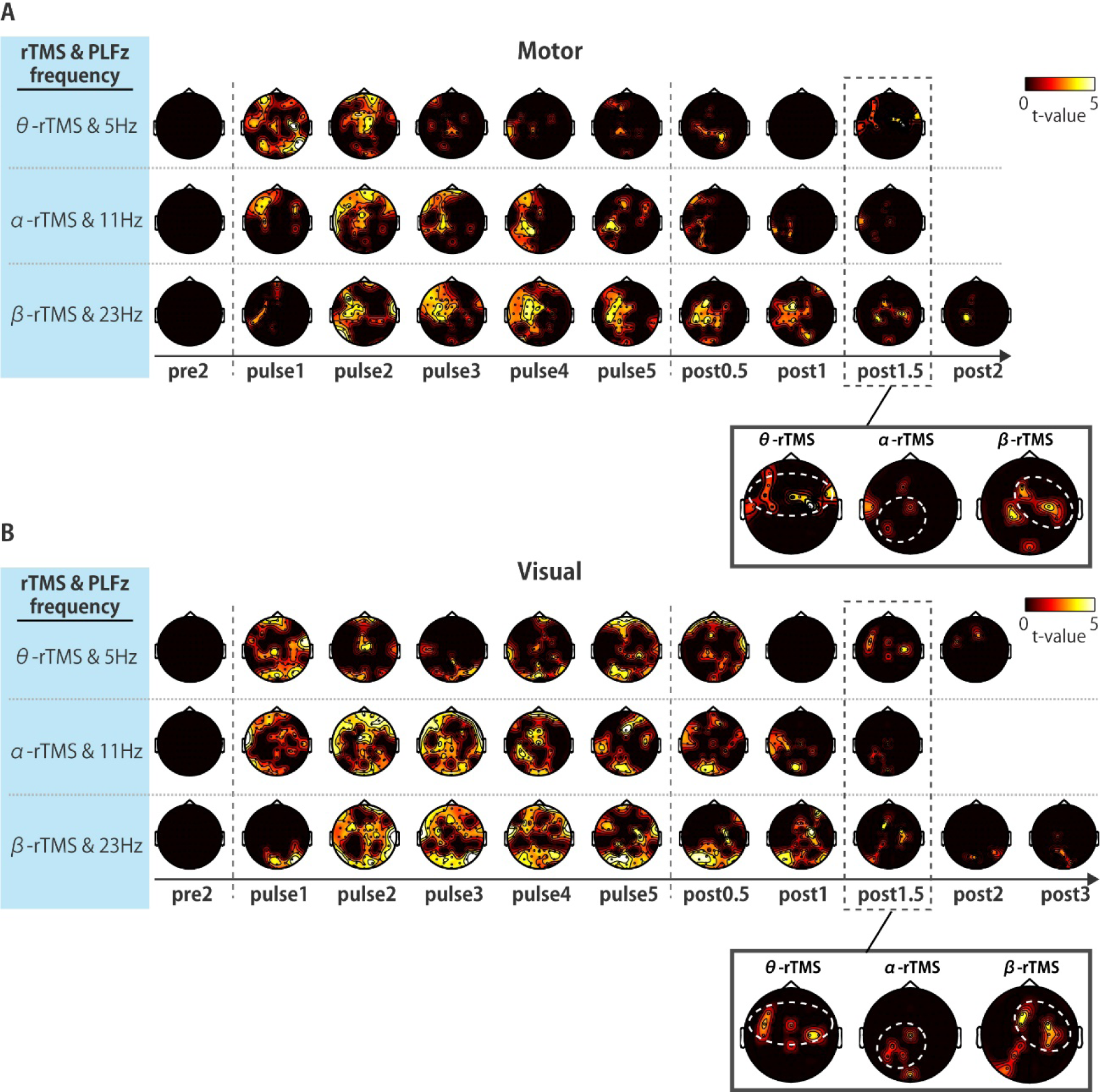
Topographic map of statistical *t*-values at the rTMS frequency. Significant clustered *t*-values from the comparison of ZPLF values between motor and sham stimulation (A), and between visual and sham stimulation (B) (*p* < 0.05). Each map corresponds to a specific frequency and a specific time point in the experiment: 2 cycles pre-stimulus, the 5 pulses in the pulse train, or 0.5 to 3 cycles after the end of the pulse train. The phase locking of theta frequency oscillations in the frontoparietal, alpha frequency in the left occipitoparietal, and beta frequency in the right motor areas are highlighted with white dotted circles in the magnified view of post1.5 cycles.

We further examined the time–frequency profile of the extent of the spatial spread of phase-locking by counting the number of channels (electrodes) with significant phase locking. For motor stimulation, phase locking at alpha-band frequencies was induced in many channels by α-rTMS (Fig. 5A and B, middle panels). On the other hand, β-rTMS induced oscillations at both alpha- and beta-band frequencies (Fig. 5A and B, bottom panels). For α-rTMS or β-rTMS over visual cortex, the induced oscillations were prominent at the stimulation frequency (Fig. 5C and D, middle and bottom panels). To quantify this frequency specificity, we averaged the number of significant channels over four cycles at each frequency (indicated by the red lines in Fig. 5A and 5C). We used the binomial test to assess whether more channels were phase locked at the stimulation frequency than at the other frequencies (e.g., for θ-rTMS, we assessed whether there were significantly more phase-locked channels at 5 Hz (theta band) than at 11 Hz or 23 Hz; see Fig. 5B and 5D). The mean number of channels with significant phase locking was greater at the stimulation frequency than at the other frequencies for α-rTMS and β-rTMS over motor (Fig. 5B) and visual cortices (Fig. 5D). We also noted that the number of channels with significant phase locking was significantly larger with visual than motor stimulation (θ-rTMS: *p* = 0.1727., α-rTMS: *p* = 0.0008, β-rTMS: *p* = 0.0019, binomial test). The fact that phase locking beyond the stimulation site was most prominent at the stimulation frequency suggests that globally coupled, frequency-specific neural oscillators in the brain networks were entrained by the rTMS.

**Figure 5.**
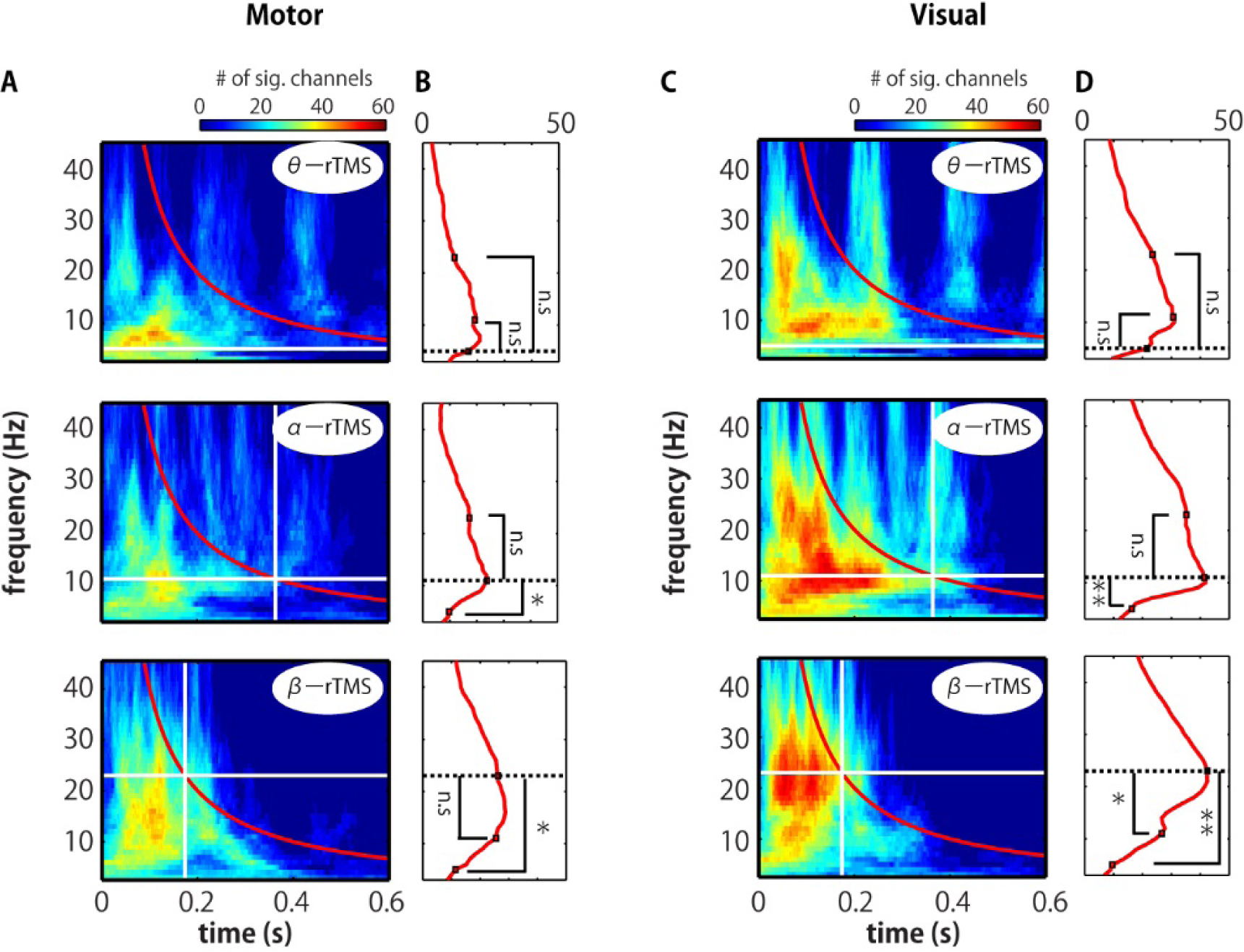
Global spread of phase entrainment. (A, C) Time–frequency representation (TFR) of the number of significant channels, determined by comparing the stimulation and sham conditions using a cluster-based permutation test. The vertical and horizontal white lines indicate the timing of the last pulse and the stimulation frequency, respectively. (B, D) Average over four cycles at each frequency (red line in panels A and C). We used the binomial test to assess whether the increase in the number of significant channels at the stimulation frequency (indicated by the dotted line) was significantly greater than the increase at other frequencies (* *p* < 0.005, ** *p* < 0.001).

## 4. Discussion

### 4.1 Efficient stimulation frequencies for the entrainment of intrinsic oscillations

Earlier behavioral studies indicated that rhythmic stimulation at a frequency that matches the frequency of physiological rhythms is most efficient at modulating behaviors via the entrainment of task-related oscillations (Klimesch et al., 2003; Sauseng et al., 2009; Romei et al., 2011). In particular, Klimesch et al. tested the effects of different frequencies of rhythmic stimulation, i.e., individual alpha frequency (IAF) + 1 Hz, IAF – 3 Hz, and 20 Hz, in a mental rotation task. Whereas rTMS with a stimulation frequency of IAF + 1 Hz was beneficial for behavior, stimulation at only 3 Hz deviation from IAF had no effect. More recently, a carefully designed TMS–EEG study by Thut et al. (2011b) showed that intrinsic alpha oscillations were entrained by rTMS at the IAF applied to the alpha source. From the perspective of nonlinear dynamical systems theory, the degree of entrainment of an oscillatory system to the rhythmic stimulation changes as a function of stimulation frequency and amplitude (Pikovsky et al., 2003). For low stimulation amplitudes, only rhythms close to the natural frequency will entrain the system. As the stimulation amplitude increases, the system becomes entrained over a wide range of stimulation frequencies. In a model evaluating the degree of entrainment by transcranial alternating current stimulation (tACS) with a comprehensive array of stimulation conditions (amplitude: 1 to 13 pA; frequency: 0 to 6 Hz;), tACS matched to the natural frequency of the network entrained the network most efficiently at the lowest amplitude (Ali et al., 2013).

According to our knowledge, this is the first study to demonstrate frequency-specific entrainment of ongoing oscillations corresponding to the natural frequency characteristic of each network module. rTMS at theta, alpha, or beta frequencies applied to either the visual or motor cortex did not entrain intrinsic oscillations at all three frequencies: entrainment was observed at specific frequencies depending on the stimulation site. Specifically, local activity during α-rTMS over visual cortex and β-rTMS over motor cortex exhibited some signatures of entrainment. We did not use IAFs, instead using a common frequency (11 Hz), yet the visual cortex was entrained. This was presumably because the stimulation amplitude was sufficiently large and the frequency selected was close to the IAF. Nevertheless, the lasting effect of entrainment was notably longer for participants whose IAF was closer to the stimulation frequency (i.e., closer to 11 Hz; see Supplemental Information; Fig. S2). It is also expected that successive phase alignment by a periodic external force of an appropriate intensity (i.e., not too strong and not too weak) and an appropriate frequency will result in a gradual increase in phase coherence across trials (Thut et al., 2011a). Given the gradual increase in ZPLF in the alpha- and beta-bands, α-rTMS and β-rTMS likely match the natural frequency of the visual and motor cortices, respectively. Our results suggest that different local network modules have their own natural frequency and that rTMS tuned to frequencies close to the natural frequency can most efficiently modulate the oscillatory dynamics.

### 4.2 Local entrainment propagates to other areas in a frequency-specific manner

We showed that rhythmic stimulation initially causes phase locking over a wide spatial area and a broad range of frequencies, but that phase locking at the stimulation frequency is predominantly propagated as the pulse train continues. Furthermore, the areas to which phase locking eventually propagated differed for each stimulation frequency. These results suggest that local entrainment may lead to global entrainment of neural oscillators with the same natural frequency in functionally coupled regions.

Prior MEG/EEG studies have demonstrated that spontaneous oscillatory activity is spatially organized in a frequency-specific manner (Mecklinger et al., 2007; Hillebrand et al., 2012; Hipp et al., 2012). Hillebrand et al. (2012) used the mean phase-lag index (PLI) to assess the mean phase synchronization between areas and found that the mean PLIs for the alpha- and beta-bands were strongest in the posterior and sensorimotor areas, respectively. The mean PLI is the average of the weight of the edges between “nodes” (areas) and assesses the importance of nodes in a network. Thus, frequency-tuned stimulation of an important node (i.e., visual and motor areas in our experiment) may drive a frequency-specific network by successive entrainment of coupled neural oscillators from local to distant brain regions in a spatiotemporally organized manner.

Furthermore, the propagated phase locking induced by rTMS over the motor cortex was less distributed over the distinct cortical regions than that induced by stimulation over the visual cortex. We anticipated that β-rTMS over the motor cortex would actually be particularly effective at inducing propagated phase locking because the functional connectivity between the motor area and other cortical areas is largely achieved via beta-band oscillations (Hillebrand et al., 2012). We speculate that the stimulation frequency used in our study (23 Hz) may be suboptimal for entraining intrinsic oscillations because the intrinsic Rolandic beta frequency over the sensory-motor strip varies among individuals in a broad range from 14 to 30 Hz. Alternatively, the functional connectivity between motor cortex and other cortices may be actually lower than that between visual cortex and other cortices, as suggested by the data from Hillebrand et al. (2012): the mean PLI in the motor area was much lower than that in the visual area. This hypothesis is plausible because the primary motor cortex is the final cortical stage for executing a motor output and hence mainly receives inputs from other brain areas, whereas the primary visual cortex is the first cortical stage to receive visual inputs and then sends the signals on to functionally connected areas.

It should be noted that it is impossible to dissociate real and spurious connectivity of two cortical regions through conventional phase synchronization analyses if the two regions are driven by a common source (Kitajo and Okazaki, 2016). However, effective connectivity indicating directional causality between two regions can be probed by local perturbations. We therefore propose that rhythmic stimulation can be used to probe causal communication between cortical regions fueled by oscillatory activity with a specific frequency.

### 4.3 Confounding factors

We speculate that increases in ZPLF during stimulation are partially caused by contamination by repeated evoked components and TMS-related artifacts. Classical sensory rhythmic stimulation elicits a steady state response (SSR), which appears as a near-sinusoidal waveform at the stimulation frequency (Regan, 1966). The SSR is partially explained by neural entrainment to the phase of the periodic stimulation (Thut et al., 2011a; Mathewson et al., 2012). Alternatively, the SSR may be caused by recurring evoked patterns associated with the rhythmic sensory stimulation (Capilla et al., 2011). If an evoked component with constant latency and polarity is superimposed on ongoing oscillations, the intertrial phase variability of the EEG signals will decrease as if the ongoing oscillation is aligned to a specific phase (Sauseng et al., 2007). Such spurious phase locking can also be produced by repeated TMS-related artifacts.

It is challenging to distinguish phenomena due to neural entrainment, recurring evoked activities, or repetition of artifacts produced by TMS. However, entrainment signatures such as cumulative phase locking and sustained effects after the end of stimulation cannot be explained by the linear sum of single evoked components (Thut et al., 2011a). Moreover, each cortical system has a specific frequency that is particularly effective for entraining the system. We also note that weak phase locking due to the auditory component of the TMS (Romei et al., 2012) potentially contaminates all conditions, including the sham stimulation condition. We always subtracted the sham condition from the actual TMS condition in all analyses. Although contamination by TMS-related artifacts remains a possibility, the neural entrainment observed in this study are more likely to have been caused by the entrainment of ongoing oscillations, rather than by repetitive evoked and/or artifact components.

### 4.4 Conclusions

rTMS can directly probe the mechanistic properties of the intact brain. This study used multiple stimulation frequencies that targeted two areas, and demonstrated for the first time that distinct oscillatory frequencies are established by the local and global mechanistic architectures of brain networks. Specifically, we showed that rTMS can entrain intrinsic local and global neural oscillations, presumably via the sequential entrainment of neural oscillators in a brain network. The entrainment was observed only when the stimulation frequency matched the natural frequency of the network. In other words, it is possible to selectively entrain synchronous networks in a frequency-specific way. The ability to directly modulate brain rhythms in a frequency and target-specific manner may provide insights into the functional roles of rhythmic activity. Moreover, this technique has clinical potential: modulation of impaired oscillations and synchrony may alleviate symptoms in patients with attention deficit hyperactivity disorder (Barry et al., 2003), Alzheimer’s disease (Prichep et al., 2006), depression (Vuga et al., 2006), obsessive–compulsive disorder (Sherlin and Congedo, 2005), schizophrenia (Lee et al., 2006), and stroke (Kawano et al., 2017).

## Acknowledgments

This study was supported by JST PRESTO, MEXT Grants-in-Aid for Scientific Research 26282169, 15H05877, and a research grant from TOYOTA Motor Corporation. We are grateful to Shun Sato, Atsushi Negishi, and Hiroyuki Okura for help in data analysis.

## Author contributions

Y.N., T.H. and K.K. designed research. Y.N., Y.O.O., Y.M., T.H. and K.K. collected the data. Y.N. Y.O.O. and K.K. analysed the data. Y.O.O. T.H. and K.K. wrote the manuscript.

## Legends

**Figure S1.**
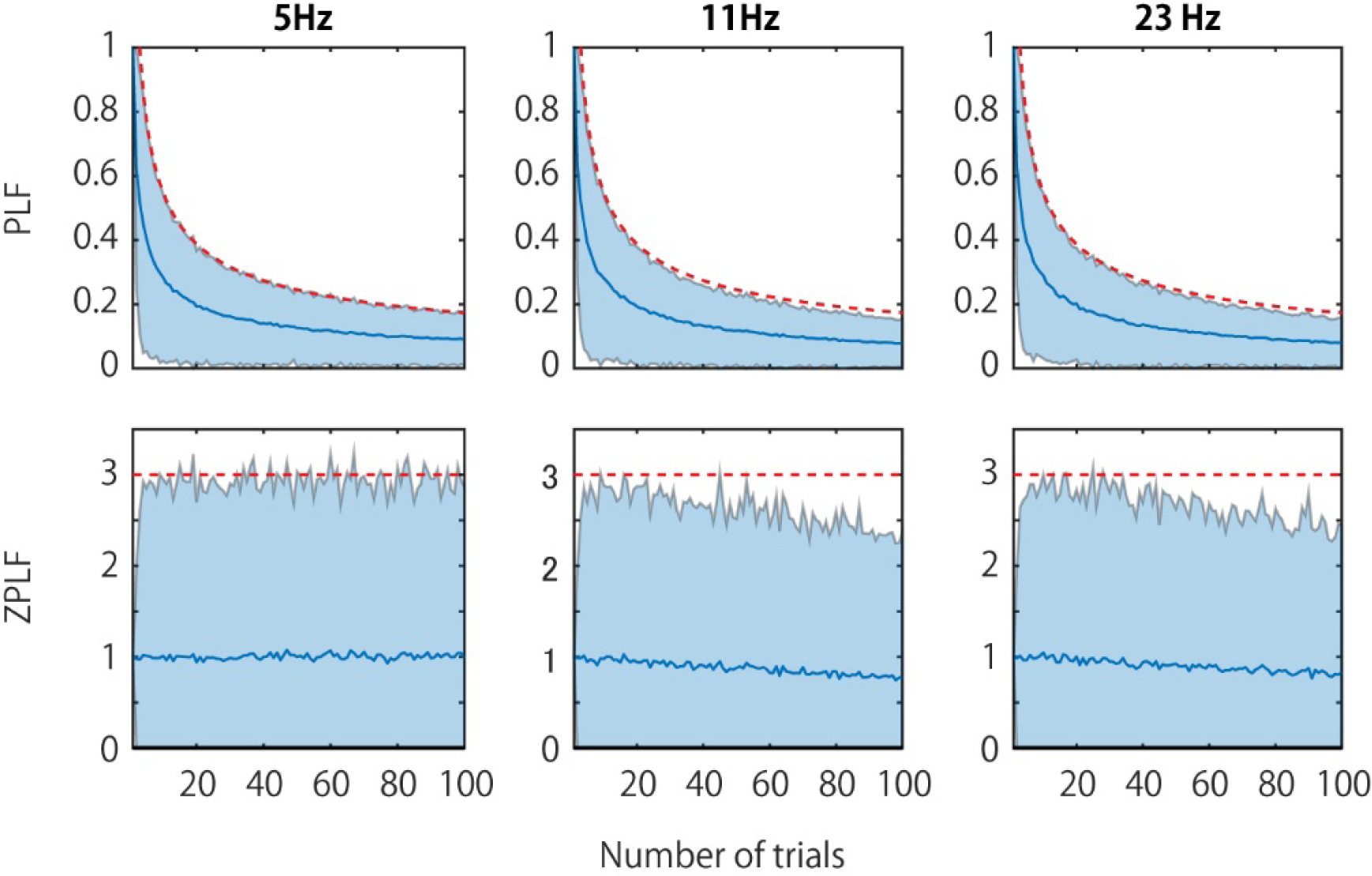
PLF (A) and ZPLF(B) as a function of the number of trials for a single subject. PLF and ZPLF at the pre-TMS time point (electrode: Cz) were computed with randomly selected trials from a trial pool of all conditions and were averaged over 1000 times iterations. The blue line and blue area are the mean and 95% confidence intervals, respectively. The red line shows the critical PLF and ZPLF values obtained by 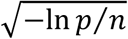 and −ln *p*, respectively, where *p* = 0.05 and *n* is the number of trials used to calculate PLF and ZPLF.

**Figure S2.**
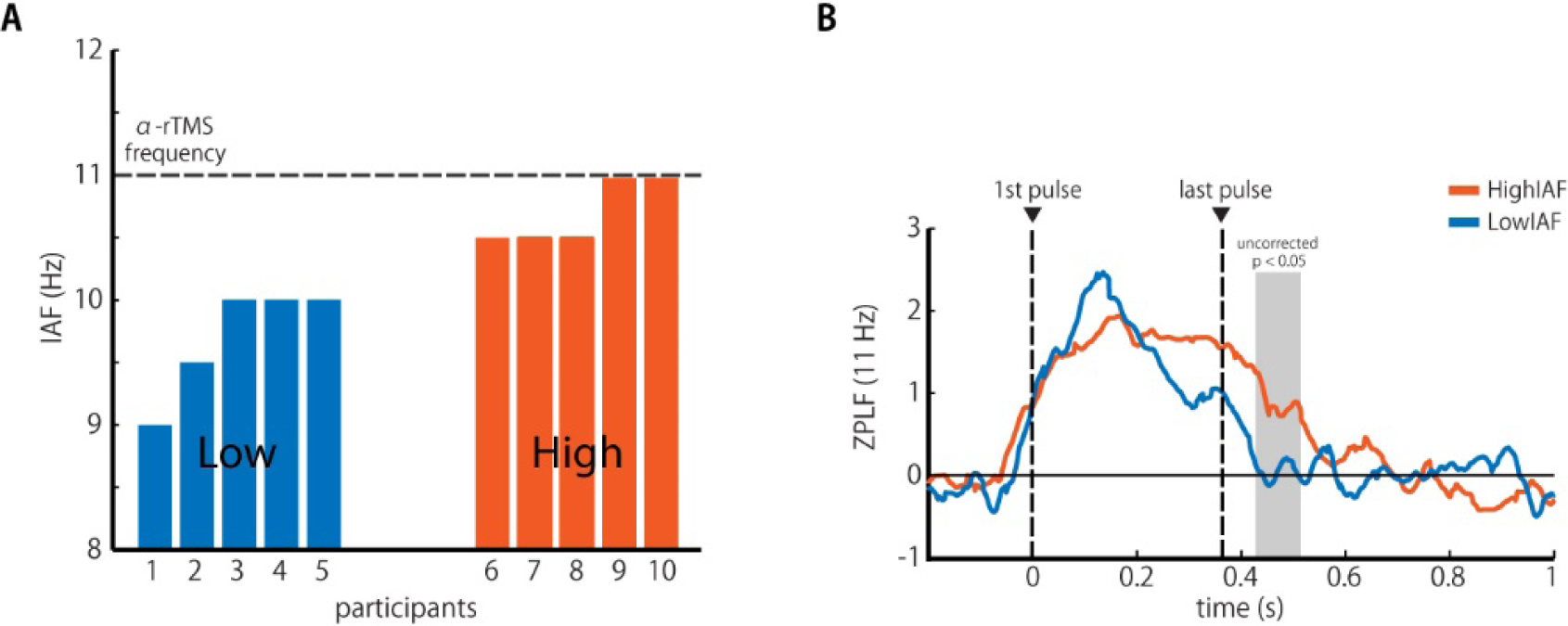
ZPLF of high-IAF and low-IAF groups. The FFT-based power spectrum was computed for 2 s data from pre-sham-stimulation epochs (Hanning window, frequency range: 2 – 45 Hz). We defined the individual alpha frequency (IAF) as the frequency with the maximum power peak at Oz, within the range from 6 to 13 Hz. (A) We separated participants into low-IAF and high-IAF groups. The stimulation frequency in the α-rTMS condition was close to the natural frequency of the high-IAF group. Four participants who did not have a prominent power peak were excluded from this IAF analysis. (B) The grand averaged ZPLF for each group for α-rTMS over visual cortex. Note that the ZPLF of each participant was standardized by the mean and SD over time points before those used for the grand average. The shaded area indicates time points during which there was a significant difference between the two groups (*p* < 0.05, uncorrected *t*-test); the lasting effect of entrainment was notably longer in the high-IAF group. These results suggest that frequency-tuned rTMS sufficiently close to the natural frequency is the most effective for entraining intrinsic brain oscillations.

